# cellNexus: Quality control, annotation, aggregation and analytical layers for the Human Cell Atlas data

**DOI:** 10.64898/2026.04.14.718336

**Authors:** Shen Mengyuan, Gao Yingnan, Liu Ning, Bhuva Dharmesh, Milton Michael, Henao Juan, Andrews Jared, Yang Edward, Zhan Chen, Liu Nora, Si S, Hutchison J William, Shakeel M Haroon, Morgan Martin, Papenfuss T Anthony, Iskander Julie, Polo M José, Mangiola Stefano

## Abstract

Large-scale single-cell atlases such as the Human Cell Atlas have transformed our understanding of human biology. Yet, the lack of a robust framework that standardises quality control, expands cellular annotation, and adds normalisation and analytical layers, limits multi-study analyses and the usefulness of this resource. Here we present cellNexus, a comprehensive resource that enhances the Human Cell Atlas collection into analysis-ready data by linking quality control layers, metadata enrichment, expression normalisation, analysis and data aggregation. These enhancements enable robust large-scale statistical modelling across studies, exemplified here by a multi-tissue map of immune cell communication during ageing. All harmonised layers are accessible via a public web interface and with R and Python APIs. By providing continuous integration with CELLxGENE releases, cellNexus transforms large cell atlas corpora into an accessible, reproducible, interoperable foundation for large-scale biological discovery and the next generation of single-cell foundation models.

## Introduction

Recent advancements in single-cell RNA sequencing have profoundly transformed our understanding of cellular heterogeneity, prompting the development of extensive single-cell atlases. Among these initiatives, the Human Cell Atlas^1^, the Human Tumor Atlas^2^ and the Human Protein Atlas^3^ aim to comprehensively map human cell types and elucidate their functions in health and disease. The Human Cell Atlas, launched in 2016, aims to create a comprehensive cellular map of the human body. Early efforts of the Human Cell Atlas consortium have provided foundational reference maps of healthy tissues, most notably the lung^4^, eye^5^, and nervous system^6,7^. The HCA consortium plans to extend its draft atlas coverage to 18 Biological Networks. Platforms such as CELLxGENE^8^, developed by the Chan Zuckerberg Initiative, have further enhanced data accessibility by offering interactive and intuitive visualisation and analysis tools for single-cell datasets. The associated Census API complements CELLxGENE by providing programmatic access to an extensive collection of harmonised single-cell datasets, streamlining data retrieval and facilitating large-scale analyses.

Despite these advances, significant barriers remain in transitioning from multi-source datasets to meaningful scientific analyses. Issues such as a lack of standardised quality control^9^, variable RNA abundance scales, diverse gene transcriptional abundance transformations, numerical challenges including negative abundance values, missing donor annotations, and inconsistent cell typing resolution limit streamlined and large-scale analyses. Furthermore, the burden of resolving such inconsistencies often falls on individual users, who may require advanced computational resources and expertise that are inaccessible to smaller laboratories or early-career researchers. As a result, the full potential of these community-contributed datasets may remain unrealised.

We developed cellNexus to address these challenges and empower researchers to conduct rigorous, reproducible analysis across heterogeneous datasets. cellNexus is a resource that bridges the gap between data and downstream analysis. This resource provides curated metadata layers that include standardised quality control metrics, harmonised and normalised gene expression values, enhanced data annotation, and aggregated cell representation for streamlined large-scale analyses. cellNexus uses continuous integration as a foundation for following the source data release cycle. Accessible through command-line and web interfaces, cellNexus empowers researchers to query, visualise, and integrate single-cell data at scale. Users can seamlessly retrieve quality-controlled metadata or counts and integrate them with the broader CELLxGENE ecosystem, including the Census API. Here, we illustrate how cellNexus directly facilitates the creation of population-level models of cell-cell communication and high-quality cellular atlases from highly curated and annotated cells. By reducing technical barriers and streamlining access to expertly curated datasets, cellNexus democratises large-scale single-cell analysis, fostering collaborative investigations and accelerating discovery.

## Results

### Overview of cellNexus

cellNexus is a quality-controlled, curated, and extensively annotated resource that facilitates and accelerates the data-to-analysis process. Currently, cellNexus harmonises over 44,016,075 cells within the CELLxGENE collection, focusing solely on primary data. A scalable, automated pipeline streams data for each sample through four quality-control layers (Figure 1A). These layers include detecting anomalous gene expression distributions, harmonising gene expression to a negative binomial-like distribution, and detecting empty droplets, doublets, and viable cells. Gene expression is normalised using counts per million or SCTransform^10^. Immune cell type annotations are consolidated via multi-reference consensus labelling; sex and ethnicity metadata are imputed using machine-learning approaches. Donor-level features such as inferred cell-cell communication are computed to support higher-order analysis. Furthermore, precalculated pseudobulk data from high-quality cells enable Human-Cell-Atlas-scale analyses, such as modelling the immune system across tissues at the population level (Figure 1B). Our continuous integration framework enables periodic updates to our resources following CELLxGENE release cycles.

**Figure 1.**
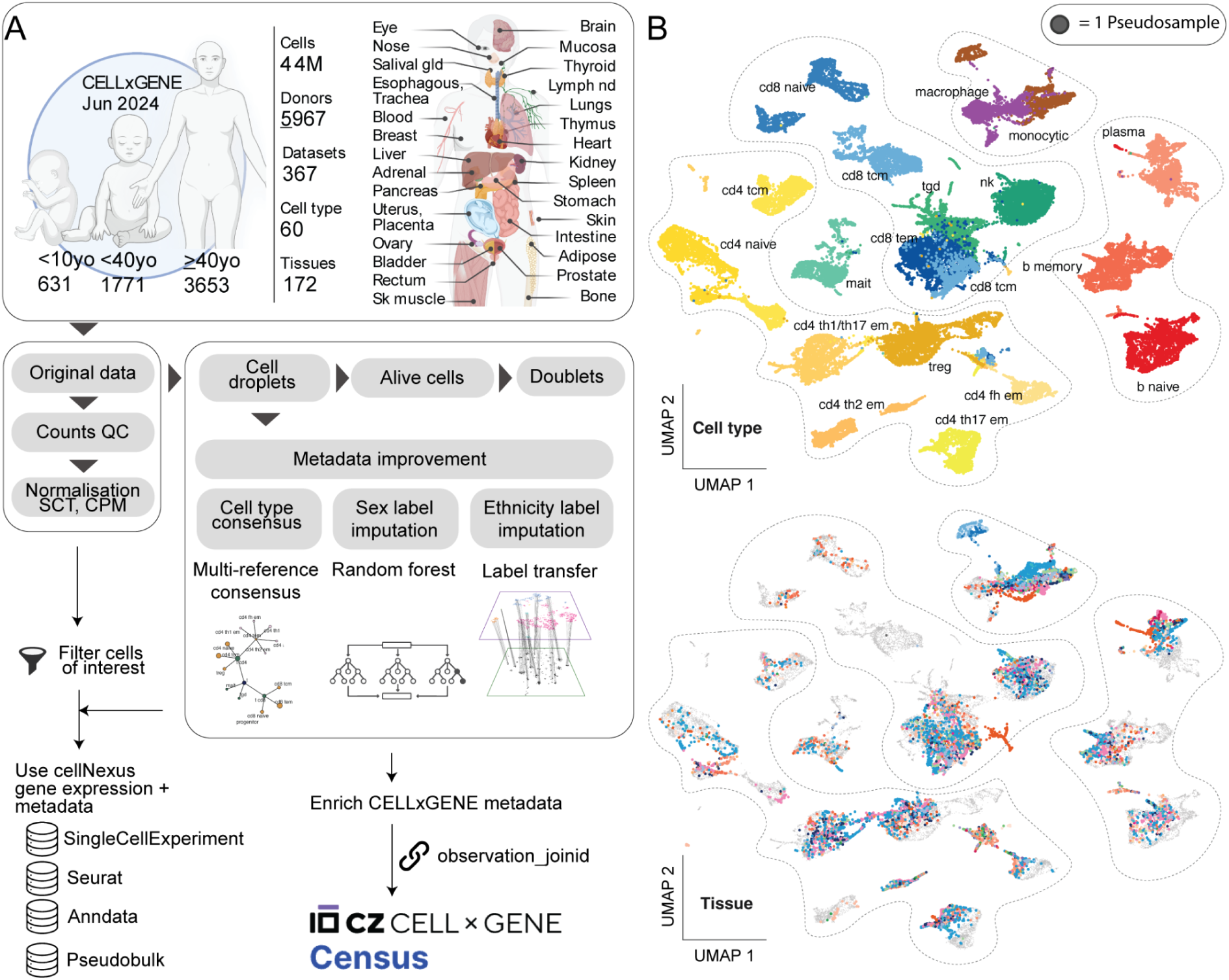
Overview of the cellNexus resource and processing pipeline. **A:** Top panel: Size of the source CELLxGENE database. Middle panel: cellNexus transforms raw single-cell data into harmonised expression matrices and enriched metadata. Raw counts undergo quality control, normalisation and cell filtering. Metadata layers are enhanced by multi-reference cell-type consensus, sex and ethnicity-label imputation, and cell-cell crosstalk inference. Bottom panel: Data are delivered in standard bioinformatic containers and cross-linked to the Census API. **B:** Pseudobulk UMAP representation of the cellNexus resource (immune cells only), coloured by cell types (top) and tissue of origin (bottom). The UMAP includes 235,289 pseudosamples across 18 immune cell type groups and 32 tissue groups.

These quality-control and enhanced metadata layers are exposed via R and Python APIs, and a web-based interface (cellnexus.org) allows users to filter cells of interest and export harmonised data in standard formats (e.g. *anndata, SingleCellExperiment*, *Seurat*, or pseudobulk). Alternatively, cellNexus annotation layers can be linked to the CELLxGENE census API via observation_joinid for individual datasets, enabling seamless enrichment of existing datasets with improved metadata and quality control. Our web interface allows all users to investigate the annotation layers, build API queries, and download data.

### The content and properties of the data

Our resource is based on cell metadata and RNA expression data for 44,016,075 cells from the Human Cell Atlas using the CELLxGENE census interface^11^ (1 July 2024 release). This release included data from 5,967 donors across 367 datasets. The majority of donors contributed blood specimens (39%), followed by respiratory (12%), cerebral lobes (8.6%), breast (4.9%), skin (3.8%), and renal (3.4%), with the rest of the tissue groups totalling 28% (Figure 2A). A total of 24 technologies and protocols were used to profile cells. Most donors (64%) were profiled using 10x Genomics 3’ technology, followed by 10x Genomics 5’ (25%). Other technologies each represented less than 3% of donors, including Plate-based (3.0%), Microwell (2.7%), and other platforms (2.1%). Healthy tissues, the primary focus of the Human Cell Atlas database, were profiled in 62% of donors. 104 diseases are represented, grouped into 12 major pathological conditions. COVID-19 (11%), Cancer (5.2%), Glioblastoma (3.7%), and Lung Adenocarcinoma (3.5%) are the most represented. While disease-linked specimens are often tissue-specific, blood is represented across nine diseases.

**Figure 2.**
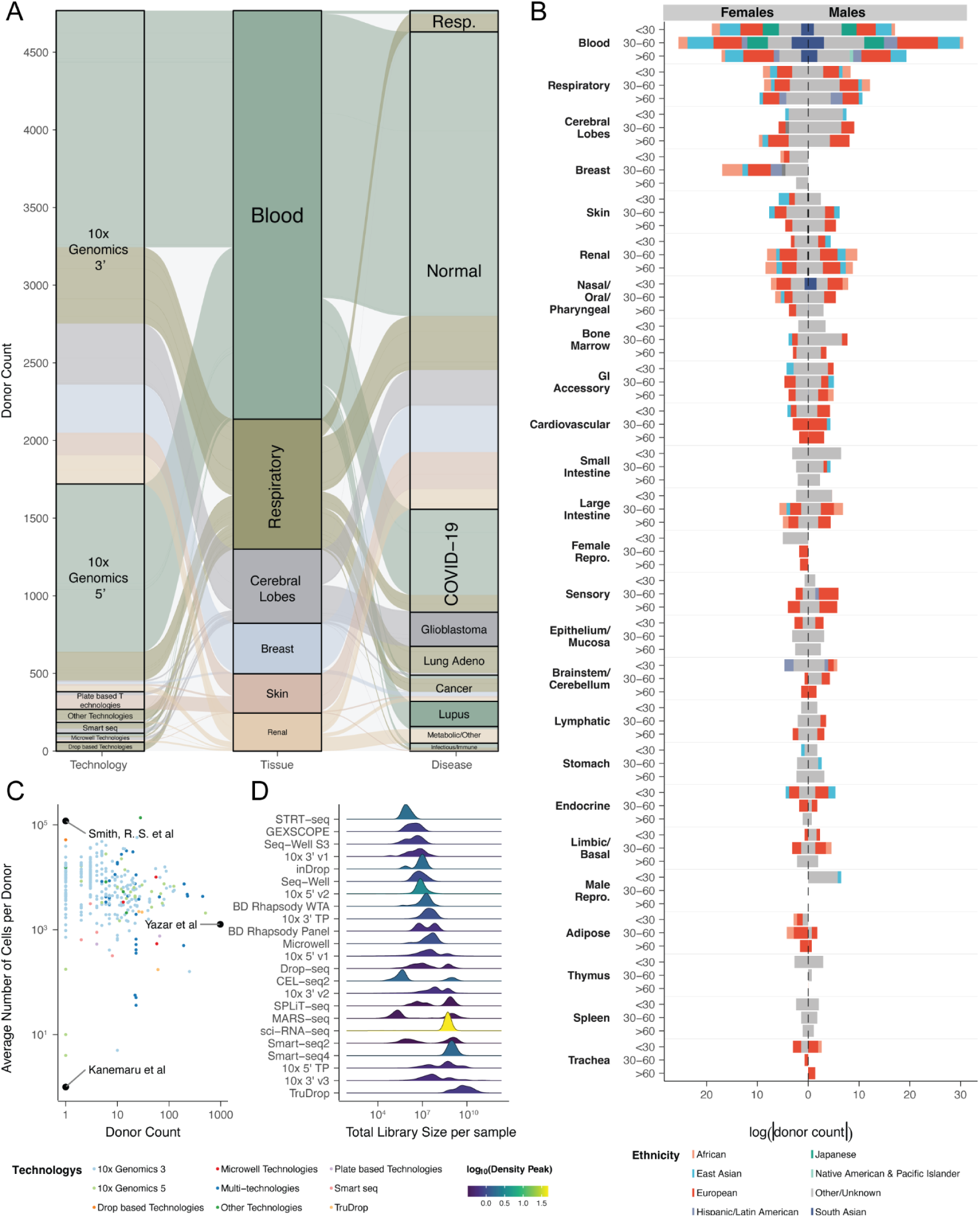
cellNexus data content. **A:** Donor counts across sequencing technologies (left) and diseases (right) for the six major tissue groups. Flows are coloured by tissue groups. **B:** Donor demographic distribution in age groups, known sex, and ethnicity groups across the top twenty-five tissue groups. From left to right horizontally, female donors are shown as negative and male donors as positive. Donors with missing sex labels are removed. **C:** Donor count and alive cell composition across datasets. Points (datasets) are coloured by sequencing technologies. Datasets using multiple technologies across samples are labelled as “Multi-technologies”. Extreme datasets are labelled with author names. **D:** Distribution of total library size per sample across sequencing technologies. The density curve is coloured by the density peak height, with brighter colours indicating higher density at that library size. Technologies are ordered by median library size (top to bottom).

The CELLxGENE data collection covers sexes and ethnicities along the ageing spectrum (Figure 2B). Males and females are represented in all tissues (excluding exclusive organs). However, bias exists, with cerebral lobes skewed towards males up to the 6th decade (54% vs 37%; 9% unknown), cardiovascular skewed towards males, especially across all age ranges(65% vs 35%), large intestine skewed towards males from the 3rd to 6th decade (62% vs 38%), adipose tissue skewed towards females across all age ranges (75% vs 25%), and skin skewed towards females below the 3rd decade (64% vs 36%). For age coverage, while blood has unbiased representation across the age spectrum, solid tissues are tightly linked to the study design and are often disease-focused. For example, the breast is limited to adults and middle-aged individuals; similarly, the prostate is biased toward seniors. In contrast, brain tissues have a broad age representation, although specific brain regions might be biased. CELLxGENE is currently biased toward Europeans (Figure 2B), representing 36% of all donors, followed by East Asians (7.9%) and Africans (2.7%). Currently, self-reported ethnicity is missing for 48% of donors.

The 367 datasets that contributed to this resource are very diverse. The smallest dataset^12^ contained one cell labelled as primary data (profiled using accepted technologies recognised by the census API), while non-primary data included 250 thousand cells (see Methods). In contrast, the largest dataset^13^ profiled 4,062,980 cells. The number of donors varies widely across datasets (Figure 2C), ranging from 1 donor in Kanemaru et al.^12^ and Smith et al.^14^, to 981 in Yazar et al.^15^. The number of cells per donor is also highly variable and is associated with the technology used. Overall, an inverse relationship is observed between the number of donors and the number of cells per donor (Figure 2C).

Technologies and protocols lead to diverse gene expression patterns and distributions. The data were often provided by researchers in transformed forms, such as log (75% of the total samples), log1p (3.7%), and log counts per million (0.20% of the total samples). The choice of transformation type is not strongly associated with the profiling technology (Supplementary Figure S1). After data distribution harmonisation (to Negative Binomial/Gamma/-like), we observed that the average per-sample library size varies by three orders of magnitude across technologies (Figure 2D). STRT-seq, GEXSCOPE, Seq-well and several 10x early versions average around 10^7^ UMIs. In contrast, Smart-seq, Sci-RNA-seq, and the most recent 10x versions average 10^9^ UMIs, with the highest being TruDrop at 10^10^ UMIs.

This comprehensive survey highlights the richness of the CELLxGENE repository and the intrinsic heterogeneity and biases arising from its scale and diverse provenance. Such complexity, spanning technologies, donors, transformations and metadata completeness, poses non-trivial challenges for integrative analyses.

### Quality control

Targeted and large-scale single-cell data analysis requires thorough, standardised quality control to increase the signal-to-noise ratio. Doing quality control is time-consuming and resource-intensive at scale. To democratise using the CELLxGENE data and facilitate the data-to-analysis process, we performed extensive quality control across cells and donors (Figure 3A). Overall, we observed diversity in quality metrics across tissue groups. For donor annotation, while tissues such as bone are particularly affected by missing sex labels, for 40% of donors, other tissues such as stomach, mucosa, bone, skin, thymus, lymph node, and spleen largely lack ethnicity labels, with up to 94% of donors affected. Similarly, cell quality metrics show diversity across tissues. Blood has the highest average empty droplet fraction across donors, at 29%, and a relatively high proportion of dead cells (1.2%) compared to other tissues. The heart, pancreas, liver, thyroid and adrenal tissue groups follow with a grand average of 13% empty droplets, while pancreas, liver, spleen and bone follow with a grand average of 0.9% dead cells. The tissue groups most affected by doublets were the salivary gland, spleen, thymus, and skeletal muscle, with percentages ranging from 3.2% to 4.6%.

**Figure 3.**
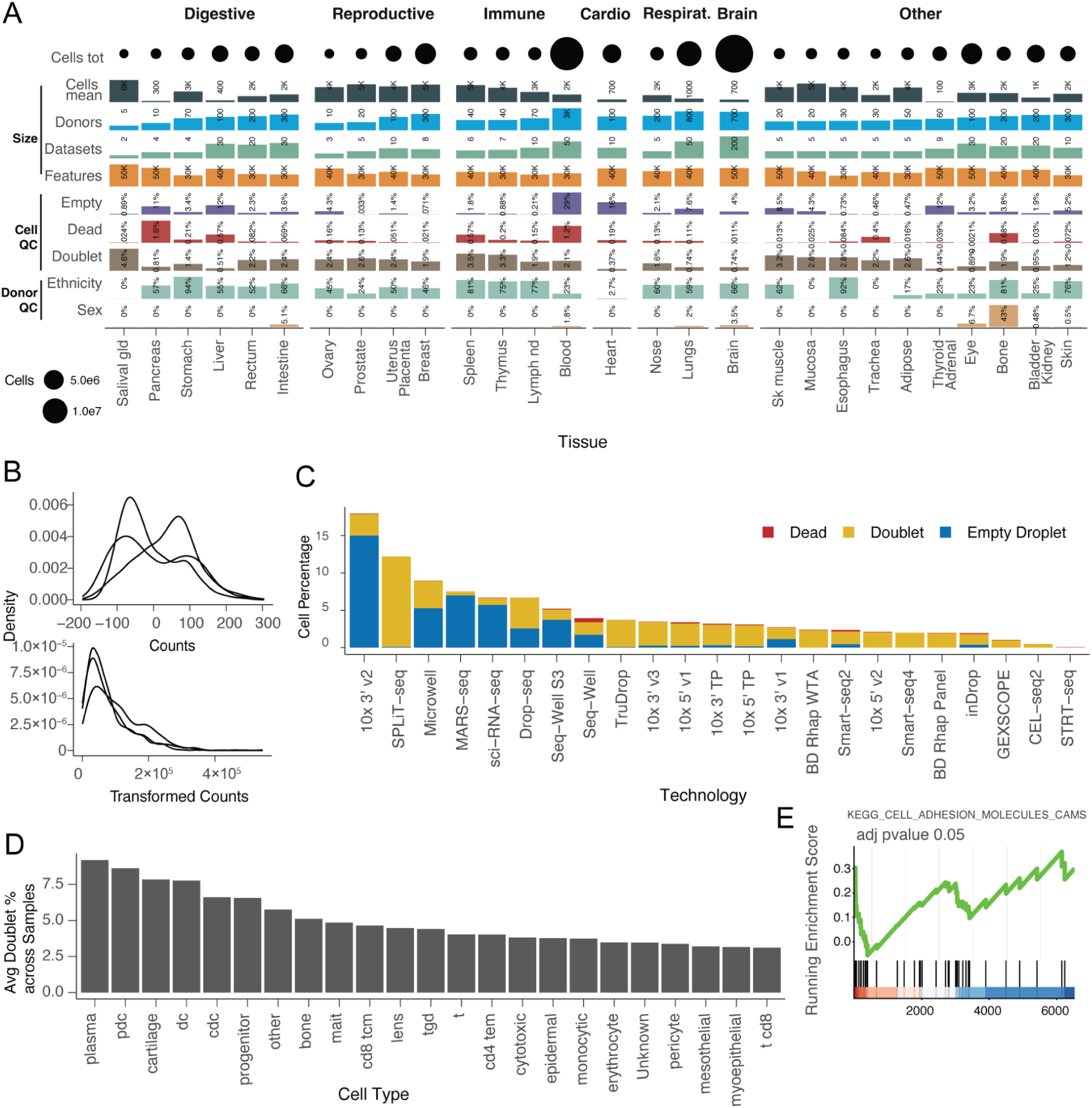
Quality control overview. **A:** Quality control metrics across tissue types, organised by tissue macrocategories. Tissues are ordered within each macrocategory by donor count. Categories: sex = proportion of donors with missing sex annotation; ethnicity = proportion of donors with unknown ethnicity annotation; doublet = mean proportion of doublets across samples; alive = mean proportion of dead cells (among non-empty); empty = mean proportion of empty droplets per sample (among alive); features = mean number of detected features per cell, averaged across samples; datasets = number of contributing datasets (log₁₀ transformed); donor = number of unique donors (log₁₀ transformed); cells mean = mean number of cells per sample. **B:** Comparison of CELLxGENE raw and cellNexus transformed counts distributions for three selected samples from three donors (IDs: 1222, 1285, 1743). **C:** Percentage of low-quality cells (empty droplets, doublets, dead cells) by sequencing technologies. **D:** Average doublet proportion for cell types across donors. **E:** Gene rank analysis.

For gene-expression quantification, we comprehensively surveyed abundance distributions across 5,967 donors, accounting for transformations and rescaling of the submitted data. We identified several distributions represented in the data, including Negative Binomial- or Gamma-distributed counts, log-normalised counts, normalised counts (e.g., scale=10,000), and log-normalised counts per million. However, we identified some non-standard distributions, such as a lower-than-zero lower boundary for 555 donors (Figure 3B; Supplementary Figure S2).

For empty-droplet identification, to enable scalability and robustness to diverse gene expression distributions, we opted for a threshold-based strategy to detect non-zero gene expression. We identified 1,765,735 empty droplets, 4.1% of the overall resource. The average proportion of empty droplets across datasets is relatively low at 1.3%. However, such proportions varied significantly across studies and profiling technologies (Figure 3C). Notably, Perez, Richard K., et al.^16^ profiled by 10x 3’ v2 exhibited the highest proportion of empty droplets at 97%. No cells from Cheng Jeffrey B et al.^17^ passed the empty-droplet threshold, with a maximum of 12 genes with nonzero expression across all cells and an average of 2.4 nonzero-expression genes per cell. In contrast, 257 highly filtered datasets do not contain empty droplets (e.g. Ren et al.^18^ and Li et al.^5^). The 10x technology with the 3’ v2 library showed the highest proportion of empty droplets (12% on average across donors), followed by sci-RNA-seq (8.7%) and Drop-seq (6.5%).

For live-cell labelling, we adopted an outlier identification strategy based on the proportion of mitochondrial gene expression. The outliers were identified per sample and cell type. We identified 31,946 (0.080% of the overall resource) of non-empty droplets as dead or damaged cells. This fraction varied widely across studies, ranging from >1.2% in two datasets^19,20^ to 0% in 165 datasets. The fraction of discarded cells was significantly associated with the profiling technologies, with Seq-Well being the most affected (0.54%; Figure 3C).

For doublet identification, scDblFinder^21^ was used for each sample. Over 90% of the donors (93%) contained doublets, whereas 11 donors exhibited doublet rates above 10% (e.g., Siletti et al.^6^). The percentage of doublets across donors averaged 2.2%, with wide variability, ranging from 0% (e.g. from Liang et al.^22^ and Reed et al.^23^) to 18% for Aldinger et al.^24^. To better understand which cell phenotypes are most prone to doublet formation, we quantified the occurrences of doublets within each cell type (Figure 3D). The average doublet proportion (averaged within each sample, then across samples) shows a high propensity for plasma cells (9.2%), followed by plasmacytoid dendritic cells (8.6%), cartilage cells (7.7%), and dendritic cells (7.6%). Beyond these top-ranked cell types, the proportion of doublets is approximately uniform across cell types (4.9% on average). RNA contamination has historically been a concern in droplet-based technologies, which can bias doublet calling, especially for transcriptionally active cell types such as plasma cells. To seek support for the validity of the doublet identification, we compared the overall transcriptome of doublets against that of singlets, within each sample in a pseudosample fashion (Figure 3E). As expected, we identified Adhesion Molecule CAMs among the significantly enriched pathways (adjusted p-value = 0.05).

This extensive quality-control survey highlights heterogeneous quality standards at the cell and donor levels. This heterogeneity underscores the need for complementary quality-control layers to convert disparate datasets into coherent, high-quality inputs for downstream analyses.

### Enhancement of donor annotations

For statistical analyses, the definition of the observational unit (i.e. sample) is fundamental. CELLxGENE does not explicitly offer a sample identifier, leaving it to the user to derive it from metadata. Considering the complexity of this dataset, this poses significant challenges in implementing statistically robust analyses. The donor identifier is the closest identifier to the observational unit. A donor may include specimens from different tissues, probed with different technologies, and at different time points (e.g., age). While reconstructing the sample identifier from the provided metadata is feasible with some knowledge of the resource, experimental batches (e.g., wells) may be hidden in the cell identifier and not otherwise provided. To streamline statistical analyses, we heuristically inferred identifiers for 41,937 samples from donor information in the metadata and cell identifiers. For example, some donors provided a significant number of cells (e.g., n=1,535,364 for H18.30.002 used across three studies^6,7,25^), which impacts both the design and the scalability of data analysis.

Our quality control (Figure 3A) revealed that 49% donors lacked at least one demographic annotation layer (age, sex, self-reported ethnicity). To address the gaps in donor annotations, we developed machine-learning strategies to impute missing labels. Self-reported ethnicity (Supplementary Table S1) is inherently complex to interpret, as self-evaluation can be biased and underreported. Self-reported ethnicity labels were first harmonised into a consistent set of broad population groups using a nationality/name-embedding framework^26^ (Supplementary Table S2). To enhance the utility of this label, we provided a low-support flag for existing labels and imputed the likely identity of missing ones. To avoid extreme tissue biases and improve the information transfer across datasets, we based our analysis on immune cells.

To distinguish ethnicity effects from other technical (i.e., dataset and technology) and biological (i.e., sex, age, disease, tissue of origin) factors, we fitted a mixed-effects linear model (brms^27^) to the donors with known labels. We identified ethnicity immune signatures, including the top 200 genes significantly associated with each ethnicity (Figure 4A). MMP8, highlighted in the African-associated set, has been linked to a functional promoter haplotype associated with preterm premature rupture of membranes in African-American neonates^28^. LGALS2, found to be up-regulated in East Asians, has been associated with a promoter variant (rs7291467) enriched in Japanese populations, which reduces transcriptional activity in vitro and is associated with myocardial infarction risk in Japanese cohorts, an effect not reproduced in British populations^29,30^. GJB2, up-regulated in European donors, is a well-established example of population-enriched variation, as the 35delG variant is much more frequent in European populations^31^. Finally, IGLV8-61, identified in the Hispanic set, lies within the immunoglobulin lambda locus, a region recently shown to harbour substantial population-level germline diversity across human groups, supporting its relevance as an ethnicity-informative immune locus^32^.

**Figure 4.**
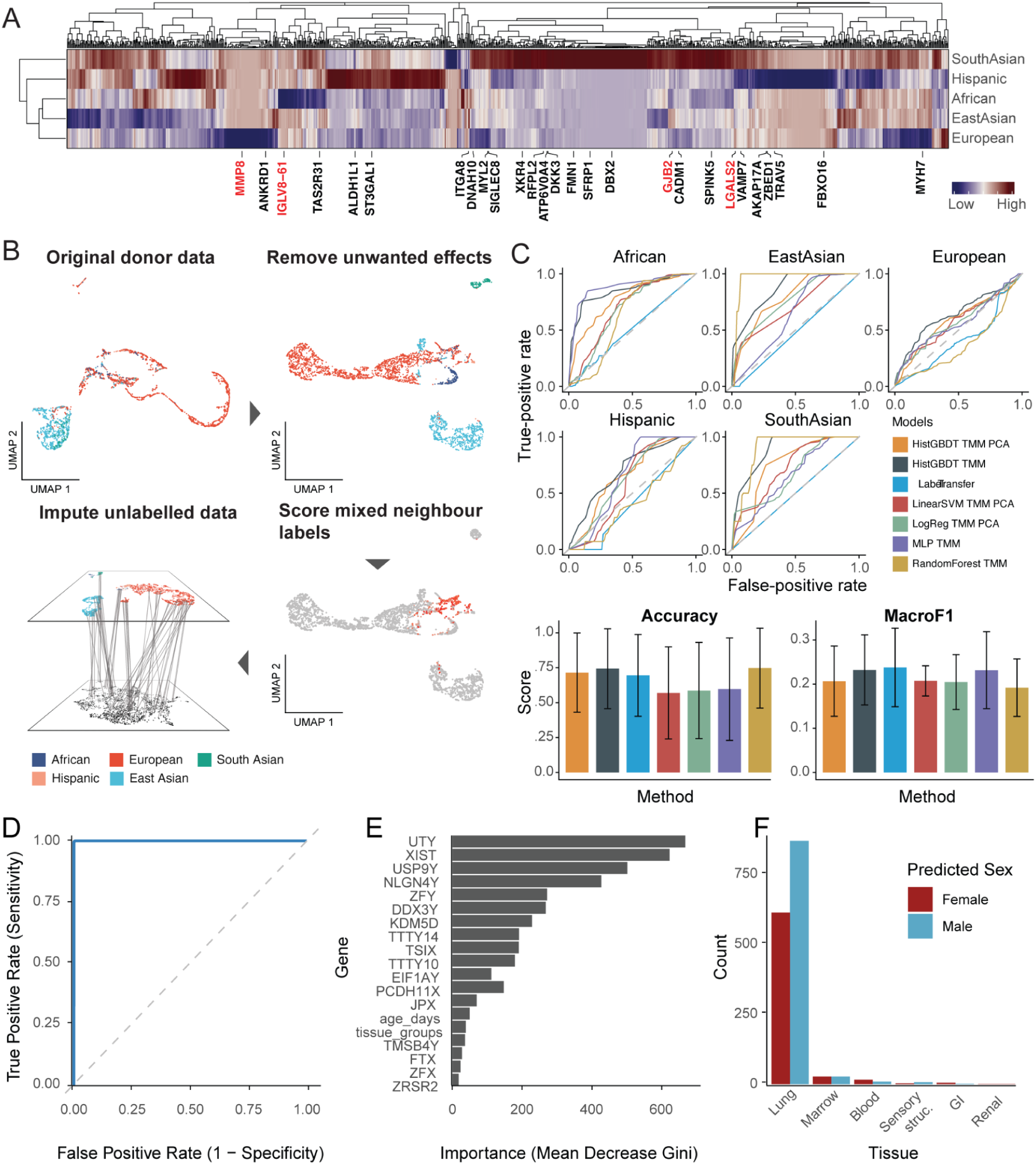
Donor annotation and label imputation. **A:** Top transcriptional markers for five major ethnic groups. **B:** Pipeline for ethnicity-label confidence scoring. From top-left to bottom-left, all non-ethnic biological and technical effects are removed, and the nearest-neighbour analysis then identifies low-confidence labels with discordant neighbours. High-confidence labels are used to transfer annotation to unannotated donors. **C:** Benchmark of missing label imputations. We benchmarked seven methods across various settings, using a leave-one-(dataset)-out cross-validation dataset to predict missing ethnicity labels from transcriptomes of pseudosamples with high-confidence labels. Top panel: Receiving operator curve (ROC); bottom panel: overall performance measured by accuracy and macro-F1 score. **D:** ROC of sex classification. **E:** Top transcriptional markers for sex. **F:** Sex prediction for missing labels across tissues.

The adjusted gene expression for the ethnicity signature was used to construct the nearest-neighbour network across donors (Figure 4B). Donors were flagged when surrounded by a discordant neighbourhood structure (using a local-graph agreement scoring metric; see Methods; Supplementary Figure S3). For example, if a European donor was positioned in an otherwise pure East Asian neighbour, that donor was flagged and excluded from the imputation process. Out of 3,259 donors, 262 were flagged as low-confidence. To impute ethnicity labels for low-confidence, missing, or unknown ethnicities (n=2,328 donors), after a stringent benchmark (see Methods), we used a histogram-based gradient-boosting classifier^33^ trained on high-confidence labels (Figure 4C; Supplementary Figure S9-S11). This model achieved the highest performance among six other methods, with median cross-validated AUC of 0.667 and 0.544, and overall accuracies of 0.713 and 0.694, under dataset-held-out and tissue-held-out settings, respectively (see Methods). The inferred labels for donors were 94% European, 3.6% East Asian, 1.5% African, 1.3% Hispanic, and 0.052% South Asian. We also identified the top markers for each ethnicity by ranking mean gene expression within each group.

To impute missing sex labels, we trained a Random Forest classifier on singscore-ranked^34^ expression from sex-chromosome genes. The model achieved high discrimination (AUC 0.982 ± 0.001, accuracy 0.971 ± 0.002) that was recapitulated on the test set (AUC 0.977, accuracy 0.970; Figure 4D). Feature importance analyses highlighted biologically coherent markers: Y-linked genes (e.g., UTY, USP9Y, NLGN4Y, ZFY, DDX3Y, KDM5D, TMSB4Y) dominated the male signal, whereas XIST was the most influential predictor of the female class (Figure 4E). Other covariates (e.g., age and tissue groups) contributed comparatively little to classification, indicating that performance is driven by robust sex-chromosome transcriptional signatures rather than dataset-specific confounders. Next, we applied the trained model to 333 donors with unknown sex labels (excluding embryonic sources). The classifier produced confident assignments and revealed tissue-wise counts of predicted female and male donors, with lung comprising the largest set, followed by marrow and blood (Figure 4F). The clear separation of predicted classes for all tissues demonstrates that the model generalises to heterogeneous biological contexts and enables consistent sex annotation of previously unlabeled donors at scale.

These analyses enriched the available donor annotations with two critical information layers, ethnicity and sex. With high prediction accuracy, the imputed labels substantially increase the proportion of data available for population-level hypothesis testing.

### Improvement of immune cell-type annotation coverage, resolution and confidence

The CELLxGENE data repository contains extensive cell annotations, providing a rich ontology to standardise cell typing within the human body. However, cell typing is particularly challenging due to the diversity of the studies and heterogeneous standards. Immune cells are ubiquitous in human tissues, and their established ontology and transcriptional markers enable harmonisation of quality standards across studies.

We used a practical, tree-based multi-source ensemble approach to improve annotation uniformity and flag cells with contrasting evidence. We integrated four pieces of evidence: the original cell annotation (Supplementary Figures S4 and S5; table available through the cellNexus API), and new labelling from Azimuth PBMC^35^, Blueprint^36^, and Monaco^37^ references (scored using SingleR^38^). We employed a tree-based voting strategy for flexible consensus identification (Figure 5A), in which the most-voted-for proximal common ancestor with an established consensus was treated as evidence when no direct consensus existed. We identified complete consensus, partial consensus with improved resolution, partial consensus with decreased resolution, and complete lack of consensus. A total of 10,508,684 cells (67%) achieved complete consensus, enriched for the subtypes macrophages (13%), NK cells (12%), and CD4 T cells (8.3%). A total of 7,099,321 immune cells had a coarse original annotation, such as B cells, T cells, and CD4 T cells. Of those, we enhanced the annotation resolution for 62% (Figure 5B). Of the 587,308 cells lacking consensus, 94% showed higher-level agreement. In contrast, 5.7% had no consensus, even at level 0, for example, when the original annotation was a T-cell subtype, and the reference-based consensus converged on a monocyte subtype.

**Figure 5.**
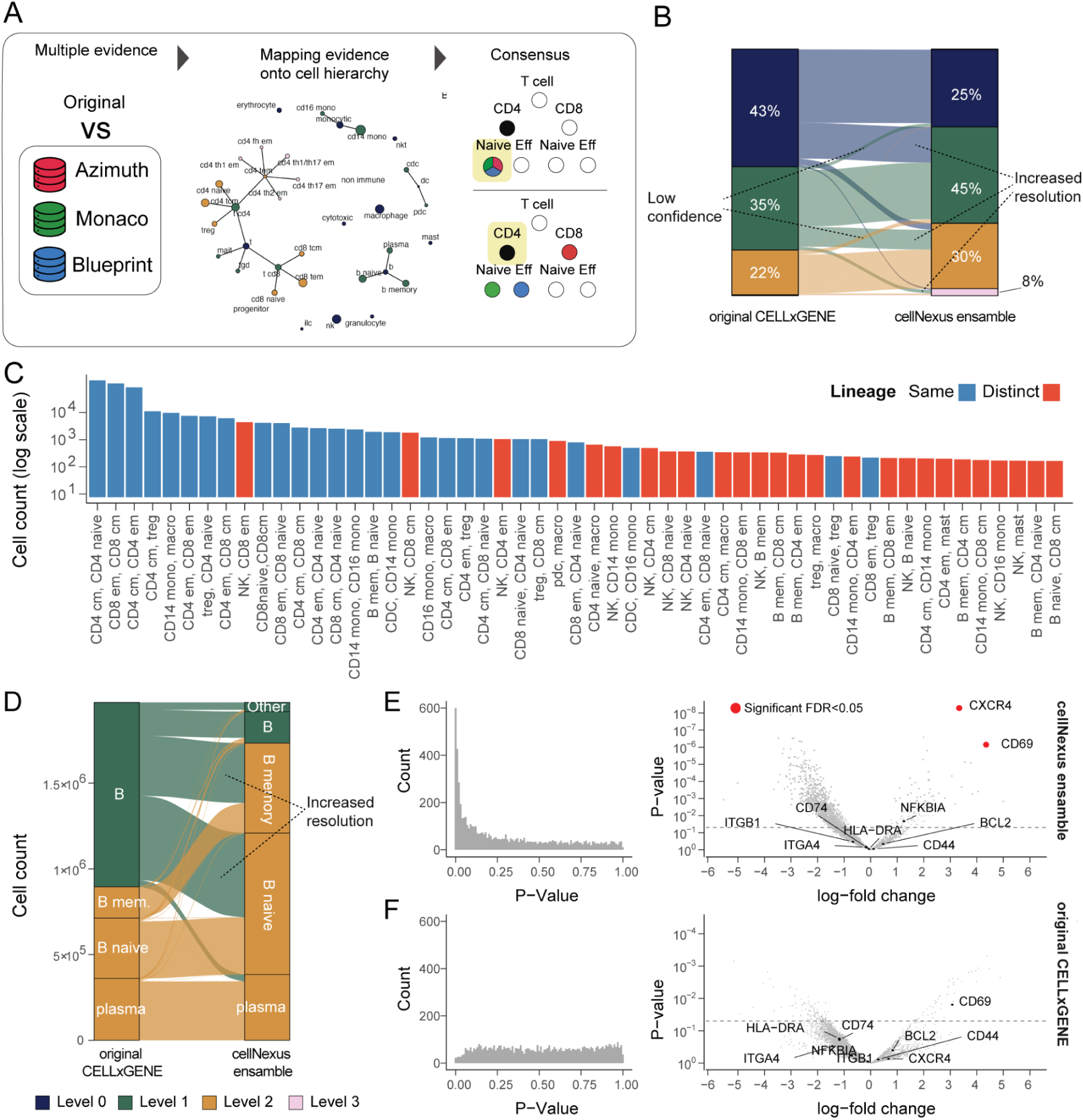
Ensemble cell-type annotation. **A:** Strategy for cell-type-hierarchy-aware ensemble cell type annotation for immune cells. The original cell-type annotation is compared with three independent annotation sources. Then, an immune-cell-type hierarchy is used to identify the optimal consensus, either at the existing, increased or decreased resolution (yellow highlight). **B:** Improvement of annotation resolution for 62% of immune cells. **C:** Rank of cell types with the most frequent discordant annotations across sources, representing challenging immune cell types to resolve transcriptionally. **D:** Type-specific improvement of annotation resolution for memory B cell, naive B cell, plasma cell, and other B cells. **E and F:** Histograms of p-values and volcano plots in differential expression analysis between blood and adipose tissue using the cellNexus ensemble annotation or the original CELLxGENE. Candidate genes whose expression is significantly enriched in adipose tissue are highlighted in red. Significance is determined by the false-discovery rate.

To better understand the causes of lack of consensus, we next sought to analyse the cell types for which the four annotations most often disagreed (Figure 5C). This analysis aimed to identify which cell types were most challenging to discern for the reference-based annotation approach. Most cell annotation disagreements represented activation states and subsets of common cell lineages (e.g., memory and naive CD4+ T cells). However, disagreement exists between more distantly related but transcriptionally convergent cell types, such as NK and memory CD8+ T cells, including 5,103 cells.

We showcase the benefit of cell-type reannotation by testing differences in gene expression between circulating and tissue-embedded (adipose) B memory cells. We increased the annotation resolution of a significant proportion of B cells (82%; Figure 5D). Compared to the original annotation, which totals 125 cells for adipose tissue across 2 samples, our consensus annotation includes 430 cells across 16 samples. Our pseudobulk analysis of the consensus annotation found, as expected, that CXCR4 and CD69, key genes for B cell tissue residency, were significantly enriched in adipose tissue (Figure 5E). In contrast, no B cell residency genes were found significantly upregulated in fat for the original cell annotation (Figure 5F).

To match the much larger Human Cell Atlas Ontology (HCAO) used in CELLxGENE, we also provide an automated alternative consensus label (Supplementary Methods; Supplementary Figure S12 and S13).

Our practical cell-type annotation pipeline, based on atlas-independent references, improved both the resolution and confidence of cell-type annotation, thereby enhancing cross-study, large-scale analyses.

### Multi-tissue landscape of ageing-related changes in immune cell communication

Providing harmonised expression matrices, refined metadata, integrated quality metrics, aggregation and analytical layers can lower the barriers to population-level analyses of biological processes. To showcase the utility of this approach, we enriched cellNexus with 30,089,091 records of cell-cell communication strength (CellChat^39^) across 241 ligand-receptor pathways, and 124,851 pseudosamples (donor-cell type combinations with sufficient count information). We use these records to build a multi-tissue map of cell-cell communication in ageing. To prioritise systemic changes, we focused on communication axes that were altered in strength across three or more tissues. The three most affected cell types with ageing (relative to the total detected pathways per cell type) were macrophages (Figure 6A), dendritic cells (Supplementary Figure S6), and CD8 T effector-memory cells (Supplementary Figure S7). Among those, macrophages uniquely showed an overall depletion of communication axes with age. To investigate the source of this bias, we ranked the most affected macrophage communication partners (either as sources or targets; Figure 6B). The cell types with the most affected communication axes (normalised to the total number of axes tested for each pair) were muscle cells, pericytes, NK cells, and endothelial cells.

**Figure 6.**
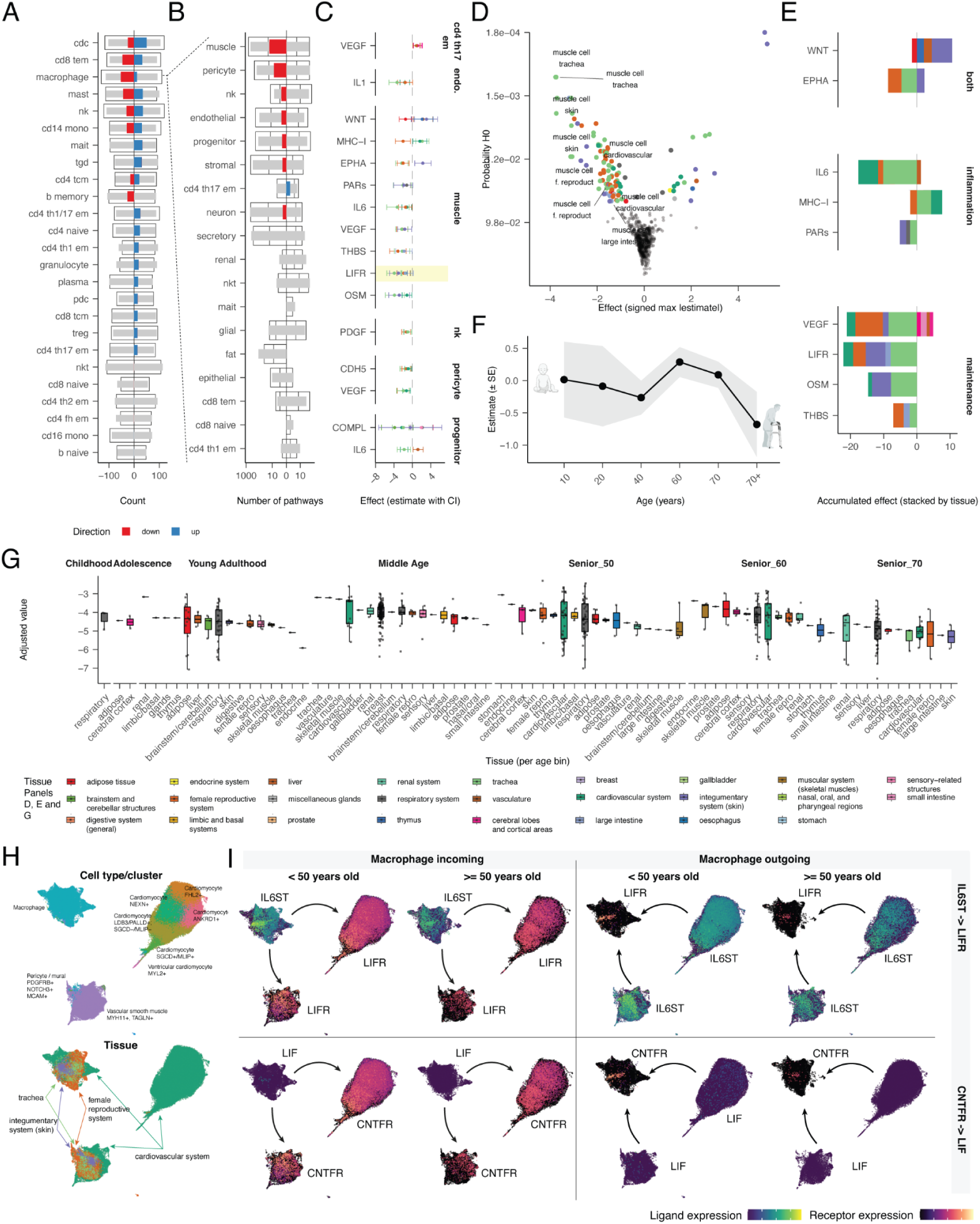
Age-associated remodelling of intercellular communication. Communication probability (log10) for each source-target-pathway-tissue combination was modelled using Bayesian mixed-effect regression (brms^27^, Gaussian). Mean model: 1 + age_bin + sex with optional covariates (assay, disease, imputed ethnicity); random intercepts for dataset and random intercepts/slopes for tissues; dispersion (sigma) had an intercept-only model. **A:** Signed counts of age-associated pathway changes per immune cell type. White bars show raw signed counts; narrow grey bars overlay sample-size-adjusted totals; coloured bars show adjusted counts restricted to pathways significant in ≥ 3 tissues. Positive values indicate increases with age; negative values indicate decreases. **B:** For the focal cell type (macrophage in downstream panels), horizontal bars summarise the number of incoming/outgoing pathway changes with partner cell types, separated by sign and ordered by the ≥ 3-tissue signal (see A). **C:** For top macrophage partners, mean effect sizes (points) with 95% credible intervals (bars) are shown per pathway and tissue. Colours encode tissues. **D:** Per-tissue pathway effects plotted against posterior null probability. Coloured points are significant after BFDR; labels highlight LIFR-axis signals. **E:** Net signed effects, stacked by tissue, for pathway groups (Supplementary Table S3). Bars sum per-tissue estimates. **F:** Population-level age_bin coefficients for macrophage→muscle LIFR across ordered bins (line = mean; ribbon = ±1 SE). **G:** Boxplots of adjusted values isolating fixed-effect age patterns for macrophage-muscle LIFR across age bins and tissues. **H:** UMAP of macrophage and muscle single cells from significant tissues. Top: clusters with labels; bottom: tissues. **I:** UMAP expression maps for LIFR-axis genes (e.g., LIF/LIFR/IL6ST; CNTF/CNTFR/LIFR), showing ligand in sources and receptor in targets, split by < 50 versus ≥ 50 years. Colour scales are per panel. The arrows indicate the ligand-to-receptor directionality.

The macrophage-muscle-cell pair had the most altered communication axes and the most depletion across tissues (Figures 6C and 6D). To better understand the leading pathways remodelled with age, we sought coordinated alteration in multiple signalling axes. The altered macrophage-muscle-cell axes of communication are part of pro-maintenance and pro-inflammatory pathways (Supplementary Table S3). Overall, we observed a shift from tissue-supportive to pro-inflammatory communication, with several canonical tissue-regenerative pathways declining with age (Figure 6E). LIFR (leukaemia inhibitory factor receptor) and its related ligand OSM (oncostatin M), both members of the IL-6 family signalling via gp130/LIFR, are implicated in muscle growth, repair, and inflammatory regulation^40^. OSM is regarded as a dual regulator of skeletal and hematopoietic tissues, with roles in regeneration and inflammation^41^. The VEGF axis, essential for angiogenesis supporting muscle perfusion and repair, is well known in regenerative biology^42^. THBS (thrombospondin) is known to act in extracellular matrix remodelling, which is required for structural integrity and tissue repair; its signalling decline would plausibly impair ECM support^43^. Conversely, pro-inflammatory signals were elevated or preserved. IL6, a hallmark cytokine in inflammaging and immune dysregulation, is frequently cited in ageing macrophage literature^44^. Persistent IL-6 signalling is associated with chronic low-grade inflammation and impaired tissue repair in aged contexts^45^. Pathways with dual or context-dependent roles, such as WNT and EPHA, exhibited more complex dynamics. WNT signalling is broadly recognised in regeneration^46^, stem cell control^47^, and fibrosis^48^. EPHA-ephrin signalling influences cell positioning, vascular patterning and inflammation in other tissues, suggesting that heterogeneous changes in EPHA may reflect spatial or niche-specific rewiring.

The LIFR communication axis was decreased in strength across the highest number of tissues (n=5). Tracking its strength across tissues by age (i.e., fixed effect; Figure 6F and 6G) shows non-monotonic trajectories, with increasing strength at 60 years old and a rapid decrease at 70 years old. This communication axis comprises several ligand-receptor pairs. These include IL6ST to LIFR and LIF to CNTFR. Mapping the expression of these genes to single cells (Figure 6H) confirms that young individuals (<50 years old) have an enriched IL6ST-LIFR pair compared with older individuals. This pattern extends to all macrophages and muscle cell subtypes, and was observed in both incoming and outgoing directions (Figure 6I). Similarly, the LIF-CNTFR is enriched in young individuals; however, the ligand or receptor expression is more limited in macrophages.

### Web server and API

To facilitate the use of the cellNexus resource, we have developed a web server and an application programming interface (API) for R and Python (Figure 7). The APIs use a fast database framework (DuckDB) that allows on-disk operations. The metadata, currently including 44,016,075 cells, can be explored, summarised and subsetted with minimal resources (Figure 7A). Metadata operations are intuitive and streamlined thanks to the tidyverse framework^49^ for R and Pandas for Python. After filtering, the quality-control and annotation layers for the cells of interest can be integrated into the source Census metadata (CELLxGENE). Also, metadata for cells of interest can be automatically packaged with normalised RNA abundance matrices into standard single-cell data containers, including anndata, SingleCellExperiment and Seurat (Figure 7B).

**Figure 7.**
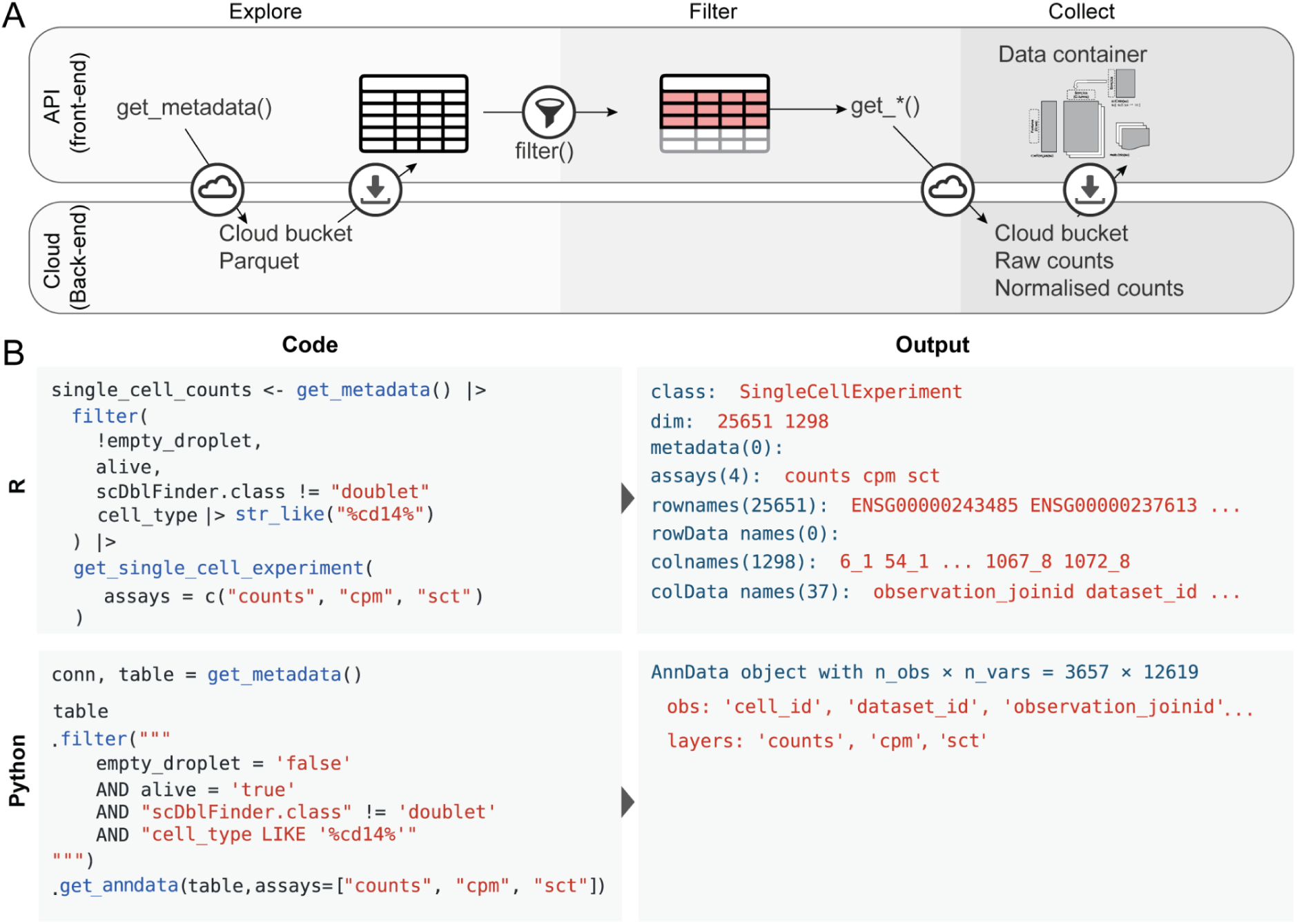
API and website design. **A:** Schematics of the API database query and data deployment. **B:** Code interface for R and Python for cell metadata and gene expression querying.

The web server (cellnexus.org) allows the broader research community to explore the cellNexus content and download the metadata and the complete pseudobulk counts (Zenodo: 17060800), including 235,289 pseudosamples and covering more than 15,000 genes. Metadata can be filtered with intuitive drop-down menus, and queries for data packaging will be returned, usable in R and Python environments.

### Continuous integration and reproducibility

To ensure a constant update of cellNexus following the CELLxGENE release cycle, we developed a continuous integration framework. An automated quality control and annotation pipeline based on R targets^50^ that identifies new data from CELLxGENE using cellxgenedp, and performs empty droplet, dead cells and doublet detection, sample annotation imputation, gene expression normalisation, pseudobulk aggregation and cell communication inference (see Methods section). This portable, reusable pipeline can be deployed on high-performance computing (GitHub: HPCell).

## Discussion

The scale and heterogeneity of public single-cell datasets have outpaced the infrastructure required to make them amenable to population-level analysis. While global initiatives such as the Human Cell Atlas have enabled data sharing at unprecedented breadth, systematic quality control, harmonised metadata, and consistent cell annotations remain missing layers, limiting reproducibility, integration, and downstream modelling. cellNexus addresses this gap by providing a curated, quality-controlled, demographically enriched, annotated, and normalised resource universe of human single-cell transcriptomes. Our pipeline implements robust cell-level QC, normalised expression, and imputation of missing metadata such as sex and ethnicity using high-accuracy machine learning. To reconcile annotation inconsistencies across studies, we introduce consensus cell typing, yielding more reliable and granular immune cell identities. These features position cellNexus as a critical tool for the next generation of single-cell foundation models.

While cellNexus represents a substantial advancement, several limitations remain. First, the resource relies on metadata and raw counts provided by primary submitters, and residual inconsistencies or incomplete information may persist despite systematic quality control and imputation. Second, although we applied robust statistical and machine learning models to infer missing attributes such as sex and ethnicity, these imputations are self-supervised and lack a ground truth; therefore, they should be considered to aid population-level statistical analyses rather than as a direct replacement for the self-reported labels. Third, the current release primarily focuses on RNA-based single-cell data and does not yet integrate other molecular layers, such as chromatin accessibility, spatial transcriptomics, or proteomics, which will be essential for fully capturing cellular states and interactions. Finally, cellNexus currently emphasises primary datasets; as more diverse populations and experimental modalities are incorporated into CELLxGENE, continuous re-evaluation of harmonisation and annotation strategies will be required to maintain accuracy and fairness across biological contexts.

By providing harmonised, analysis-ready data across millions of cells, cellNexus contributes to the democratisation of single-cell research. Without requiring users to individually download, linearise, normalise, and annotate thousands of datasets, our resource opens new avenues for comparative and gene-centric analyses directly across the Human Cell Atlas. This resource lowers the barrier to entry for groups without access to large-scale computing infrastructure, enabling rigorous hypothesis testing at tissue, cell-type, or gene resolution. Furthermore, by standardising cell- and sample-level metadata, cellNexus empowers population-scale meta-analyses and molecular association studies that were previously out of reach for most users. This shift, from data aggregation to data exploration, enables a broader, more diverse community to engage in systems-level biology and accelerate scientific discovery. cellNexus has been designed to be extensible. We plan to enhance this resource with extended cell-type consensus, beyond the immune system, and based on alternative algorithms^51^. Also, we plan to integrate multimodal, multi-species and spatial data, including the curation of disease-specific compendia^2^.

## Code availability

The code to reproduce the figures is available at github.com/MangiolaLaboratory/cellNexus_article. The R API is available on Bioconductor and at github.com/MangiolaLaboratory/cellNexus. The Python API is available on PyPI pypi.org/project/cellnexuspy/0.1.0/ and at github.com/MangiolaLaboratory/cellNexusPy. The continuous integration pipeline is available at github.com/MangiolaLaboratory/HPCell.

## Data Availability

Cell metadata is queryable via the cellNexus API (www.cellnexus.org). The pseudobulk representation is available at https://zenodo.org/records/17060800. The cell typing data is available at https://zenodo.org/records/19909778. The cell communication estimation is available at https://zenodo.org/records/19600694. The website of cellNexus is cellnexus.org, where data querying is documented.

## Author information

### Authors Contributions

SM proposed and designed the study. MS, MM, JA, JH and EY developed the computational infrastructure. MS performed quality control and summary analyses. DB and YG performed cell typing analyses. NiL performed ethnicity imputation. SM performed the communication analyses and sex inference. MS, SS and WH developed the continuous integration (HPCell). SM, JP, ATM, MM, JI, HS, CZ and NoL contributed to study design and supervision. All authors contributed to writing the manuscript.

## Acknowledgements

We thank the Human Cell Atlas Consortium for the CELLxGENE data source. We thank the Chan Zuckerberg Initiative for formatting and organising the data used in this study. We thank all individuals who donated tissues. We acknowledge all members of WEHI’s ITS and Research Computing Platform, and the members of Adelaide University Phoenix HPC for their support in computing. We thank the Stan community for their constant support. This research was supported by the Nectar Research Cloud, a collaborative Australian research platform supported by the NCRIS-funded Australian Research Data Commons (ARDC). We acknowledge the WEHI Milton HPC team that supported most of the computation. The Novo Nordisk Foundation Centre for Stem Cell Medicine, reNEW, is supported by a Novo Nordisk Foundation grant (NNF21CC0073729). We thank Prof Matthew Ritchie (WEHI) for his continuous support. We also thank Dr Milica Ng, Dr Aarathi Sugathan and Cynthia Liu for their long-standing support.

## Methods

### Data preprocessing

To harmonise gene expression matrices across heterogeneous datasets, we implemented an automated procedure to infer the transformation state of each submitted expression matrix. This step was required because matrices in CELLxGENE are contributed by multiple studies and may be stored as raw counts, normalised linear-scale abundances, log-transformed values, log1p-transformed values, or non-standard transformed quantities. When a transformation state was explicitly available from the source metadata, this annotation was used directly. For samples lacking this information, the transformation state was inferred from matrix-level summary statistics. For each sample, we computed a compact set of diagnostic features from the submitted expression matrix: whether any values were negative, whether the maximum value exceeded 20, whether all non-zero values were integer-valued, and whether non-integer continuous values were present. These features were used in a rule-based classifier designed to distinguish common transformation states while remaining robust to floating-point storage of count-like data. Matrices with non-negative, bounded, non-integer values were classified as log1p-transformed. Matrices with non-negative values and maximum expression greater than 20 were classified as raw or linear-scale abundances, irrespective of whether values were integer-valued or floating point. Matrices containing negative values but bounded within the expected transformed range were classified as log-transformed. Matrices containing both negative values and values greater than 20 were treated as non-standard linear-scale inputs and processed as raw-scale matrices for downstream harmonisation. To reduce the influence of extreme values during inverse transformation, expression values were bounded before linearisation. An upper bound of 10 was applied to log-transformed matrices with negative and bounded values, whereas an upper bound of 20 was applied to all other inferred transformation states. The inferred transformation state was then used to select the corresponding inverse operation: raw or linear-scale matrices were left unchanged, log1p-transformed matrices were linearised using the inverse log1p transformation, and log-transformed matrices were linearised using the exponential transformation. The resulting linearised matrices were subsequently passed to the centring, truncation and scaling steps used to generate harmonised count-like expression values for downstream quality control, normalisation and aggregation.

To ensure computational stability and biological interpretability across studies, we implemented a standardised transformation pipeline (GitHub: HPCell). Raw counts data (𝐶) were first scaled by a factor *S* to prevent computational overflow during downstream processing. The scaled data were then reverse processed, reversing the normalisation function (𝑓) declared by the authors at the time of submission (e.g., log-normalised counts, normalised counts with scale=10,000, and log-normalised counts per million). To maintain biological interpretability, transformed counts were centred around their most probable value (mode), calculated using kernel density estimation on a subsample of 5,000 cells for computational efficiency, and negative counts were capped at zero. Finally, cells with zero total expression were removed. The transformation process follows these steps:

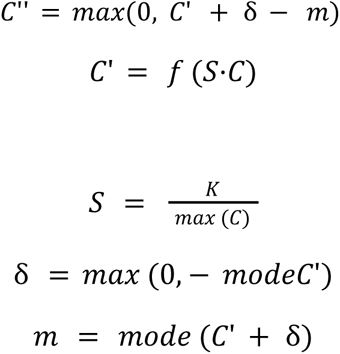

Where 𝐶 is the raw count matrix; 𝑓 is the transformation function (e.g., *exp*, *expm1, identity*); *S* is a scalar multiplier, ensuring the scaled input remains within a numerically stable range; 𝐾 is the count upper bound, set to 10 for log-distributed samples exhibiting negative values with bounded range, and to 20 for all other samples; 𝐶’ represents the scaled and transformed counts; δ is a bias-correction term that shifts the distribution to ensure non-negativity; if the mode of 𝐶’ is negative, it compensates for it by its absolute value; 𝑚 is the mode of the shifted counts; and 𝐶’’ represents the final transformed counts, centred around the mode and truncated to zero.

### Quality control

The summary statistics were based on accepted technologies from the Census API (chanzuckerberg.github.io), according to the CZ CELLxGENE Discover Census Schema: https://chanzuckerberg.github.io/cellxgene-census/cellxgene_census_schema.html.

To identify and remove empty droplets, a feature-based thresholding approach was implemented. Empty droplets were classified based on the number of expressed genes (excluding mitochondrial and ribosomal genes). Droplets with fewer than two hundred expressed genes were classified as empty. The thresholds were chosen to distinguish between cells containing biological material and empty droplets that may contain ambient RNA but lack sufficient gene expression to represent viable cells. For datasets generated by the BD Rhapsody Targeted mRNA platform, which profiles 460 genes, the threshold was adjusted proportionally and set at eleven.

The live cell identification process aimed to distinguish viable cells from dead cells based on gene expression patterns. First, the percentage of mitochondrial gene expression was calculated, as dead cells typically exhibit elevated mitochondrial gene expression due to compromised cell membrane integrity. Then, outliers were identified by high mitochondrial content using statistical outlier detection methods^52^, and labelled as dead. Cells were annotated as alive if they did not exhibit high mitochondrial expression.

scDblFinder^21^ was used to identify doublet cells using the default parameters. The algorithm was run without any predefined cell-type clustering constraints. We performed a systematic differential expression analysis to characterise the transcriptional differences between singlet and doublet cells. The analysis was designed to identify genes and pathways consistently upregulated or downregulated in doublet cells compared with singlet cells across multiple datasets. Cell quality filtering was applied to ensure only high-quality cells were included in the analysis. Specifically, only cells that were not empty droplets, were alive, and had a feature count of at least 8,000 were retained. Cells were classified as either “singlet” or “doublet” using scDblFinder^21^, and only samples with a doublet proportion greater than 5% were selected for analysis to ensure sufficient doublet representation. To maintain statistical power, only datasets containing at least 10 samples were included, and only samples containing both singlet and doublet cells were retained for comparative analysis. Single cells were aggregated into pseudobulk samples by combining cells within each sample and the doublet classification group^52^. This pseudobulk approach reduces technical noise while preserving the biological signal. It enables the use of well-established bulk RNA-seq differential expression methods that are more robust than single-cell specific approaches. Analyses were performed through the tidybulk^53^ and tidyomics^54^ interfaces. Gene selection was performed to ensure cross-sample comparability by retaining only genes present across all samples.

Abundance filtering was applied to remove genes with fewer than 20 counts per million in at least one condition, and TMMwsp (Trimmed Mean of M-values) normalisation^55^ was applied to account for compositional differences between samples. Differential expression was tested using edgeR’s robust likelihood ratio test^56^ with a design formula of ‘∼ scDblFinder.class + sample_id’ to account for both the doublet status and sample-specific effects. Genes with absolute log2 fold change greater than 1 were prioritised for further analysis, and p-values were adjusted using standard multiple testing correction methods to control for false discovery rate. Gene set enrichment analysis^57^ (GSEA) was performed to identify biological pathways associated with doublet status. Genes were ranked by log2 fold change from the differential expression analysis, and MSigDB^58^ C2 curated gene sets were used for pathway analysis. Ensembl gene IDs were mapped to Entrez IDs to enable pathway analysis, and the analysis was performed specifically for Homo sapiens.

### Sex label quality control and imputation

We designed an end-to-end workflow to infer biological sex from large-scale transcriptomic data. We aggregated gene-level counts to sample-level pseudobulk, annotated gene models with genomic coordinates, extracted and transformed sex-chromosome features, and fit a random forest classifier under cross-validation. Performance is summarised by internal resampling and a held-out validation cohort built from sex-specific tissues (e.g., breast and prostate).

To focus the feature space on loci that are informative for sex determination, we annotate gene coordinates and derive sex-chromosome features. We mapped gene identifiers to chromosomes using org.Hs.eg.db (v3.23.1) and kept only genes on chromosomes X and Y. We converted counts to a rank-based^34^ representation. The rank transform standardises marginal distributions across samples and studies while preserving order information, thereby improving robustness to library size and compositional shifts.

To limit confounding by developmental context, we remove likely embryonic/fetal samples where possible, retaining adult and adult-like profiles (e.g., ‘age_days ≥ 365’ or cases with missing age but without fetal/placental keywords in titles). We categorise tissues as male-specific, female-specific, or other based on prior biological knowledge (for example, the prostate as male-specific and the breast/ovary/female reproductive system as female-specific). We evaluated performance using 5-fold cross-validation on 2,726 training donors (70%) and a held-out test set of 779 donors (20%). The validation cohort (10%) comprises only sex-specific tissues and is never seen during training; the training cohort contains the remaining tissues. When a study lacks sex-specific tissue samples, we create a fallback validation cohort by randomly sampling sex-balanced profiles. For labelled data not assigned to validation, we create a stratified train/test split that preserves sex-class balance in both partitions.

Model training uses tidymodels (https://github.com/tidymodels/). When present, predictions are constructed by column-binding the sex-chromosome expression representation with available metadata covariates, such as tissue group or batch. The recipe imputes missing numeric predictors via the median, removes zero-variance predictors, normalises numeric predictors to mitigate scale differences, and prunes highly correlated predictors at a 0.95 threshold to reduce redundancy and stabilise the subsequent learner. We train a ‘randomForest’ classifier, chosen for its resilience to noisy predictors, invariance to monotone transformations, and capacity to model nonlinear interactions. During tuning, we evaluate accuracy, area under the ROC curve, sensitivity, and specificity, then refit the best workflow on the full training set.

We quantify robustness and generalisation at multiple levels. First, cross-validation metrics summarise the central tendency and variability under resampling. Second, test-set metrics from the internal split measure performance on held-out labelled samples from the same tissue distribution. We compute overall metrics on this cohort and break out accuracy by tissue. To aid interpretation, we extract feature importance. We used the trained model to impute unlabeled samples.

### Ethnicity confidence scoring and imputation

This analysis is strictly of self-reported ethnicity, as provided by CELLxGENE (Supplementary Table S1). Therefore, it should not be intended as a genetic ancestry analysis. Self-reported ethnicity labels were first mapped to a standard ethnicity grouping (Supplementary Table S2). Ethnicity groups with fewer than 20 representatives were excluded from inference, including those from the Oceanian and the Americas. Additionally, samples with self-reported ethnicity as “Asian” were excluded to minimise confusion between East and South Asian ethnicity groups. To identify gene expression-based ethnicity markers, we performed a pseudobulk analysis of the immune system to reduce tissue-specific effects. We used brms^27^ with the Hamiltonian Monte Carlo algorithm to fit a zero-inflated negative binomial of the pseudocounts. Priors were weakly informative and adapted to the scale of the interaction weights, with Student-t priors for intercepts and fixed effects, exponential priors for standard deviations, and LKJ priors for correlation matrices. Datasets were modelled with a random intercept, and ethnicity (together with sex and age) was modelled as a random effect against tissue group label. The model formula used to model gene expression was: ∼ offset + ethnicity_group + age + sex + technology_group + disease_group + (1 | dataset) + (ethnicity_group + age + sex | tissue). The offset was calculated as the log scaling factor from the TMMwsp (Trimmed Mean of M-values) normalisation^55^. Using the fitted model,^27^ from the data, while preserving ethnicity, we obtain adjusted pseudocounts. Ethnicity-associated marker genes were defined for each ethnicity by retaining genes with positive ethnicity-specific model estimates and ranking them by the positive lower bound of the corresponding confidence interval, followed by the effect size estimate; the top 200 genes per ethnicity were selected, and their union was used for downstream analysis. Using adjusted pseudocounts for the union of ethnicity-associated marker genes, we projected all donors into 30-dimensional principal component (PC) space. We reassessed the ethnicity labels of donors using a local-graph agreement score, and then imputed labels for the unknown/low-confidence donors using a histogram-based gradient-boosting classifier. For each donor 𝑖, we identified its 𝑘_𝑙𝑜𝑐𝑎𝑙_= 15 nearest neighbours in the PCA space and computed the local agreement α = 𝑝(𝐿_𝑖_), where 𝑝_𝑖_(𝐿_𝑖_) is the fraction of neighbours with the same ethnicity label as the donor 𝑖. We further defined a local margin 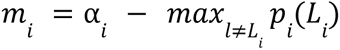, representing separation from the strongest competing label, and calculated the confidence score as

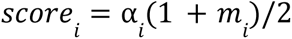

Donors with confidence scores below 0.55 were classified as low-confidence labels.

To test the accuracy of the outputs, we performed a quality control on four donors with discordant neighbours (Supplementary Figure S3). We retrieved each donor’s nearest neighbours (k = 10) from the PCA space kNN graph. We overlaid the target donor and its neighbours on the global UMAP, colouring by ethnicity. In all four cases, neighbours were mostly concordant with the target donor, consistent with the high local agreement observed in our scoring step. Therefore, the apparent offsets in UMAP likely reflect non-linear embedding distortions rather than mislabelling.

To impute ethnicity labels for unknown and low-confidence donors, we first examined tissue-by-ethnicity distributions to assess potential confounding (Supplementary Figure S8).

Because study-level and tissue-level effects were expected to be a major source of batch structure, we evaluated ethnicity-imputation models using both dataset-held-out and tissue-held-out cross-validation. We trained classifiers on the high-confidence donors using the same PC features and compared logistic regression, random forest, support vector machine (SVM), Seurat label transfer, histogram-based gradient boosting, and a multilayer perceptron (MLP). We assessed performance using accuracy, macro-F1, and one-vs-rest ROC analysis for each ethnicity (Figure 4C and Supplementary Figures S9-S11). The final model was a histogram-based gradient-boosting classifier trained on log-transformed, TMM-normalised expression of the selected marker genes after 30-dimensional principal component analysis, as this configuration showed the most robust performance across evaluation settings.

### Cell typing

We developed a data-driven, knowledge-graph-aware ensemble framework to harmonise cell-type annotations across large-scale single-cell datasets. The approach integrates predictions from multiple widely used methods into a unified immune-cell ontology, including Azimuth PBMC^59^, Blueprint^36^ and Monaco^38^ (scored using SingleR^38^), and the original CELLxGENE annotations. A directed immune hierarchy serves as a knowledge graph to reconcile disagreements at fine resolution, and the framework selects the deepest consistent node where support converges. We implemented two complementary consensus strategies: a conservative rule based on the lowest common ancestor and a weighted vote propagation model that leverages method- and prediction-level confidence.

Input data and harmonisation followed a reproducible workflow. We harmonised predictions using a shared immune-cell dictionary from Azimuth PBMC^59^, Blueprint^36^ and Monaco^38^, and the original CELLxGENE labels. All harmonised labels were embedded in a directed immune hierarchy represented as an adjacency matrix, which provided the backbone for integration by enabling ancestor and descendant queries and graph distances.

We computed a consensus label from the harmonised method predictions for each cell using the knowledge graph to guide conflict resolution. Predictions that could not be mapped into the dictionary were set to missing before consensus. The overarching principle was to prefer the deepest possible node in the immune hierarchy when predictions converged, and to back off to parent terms when predictions diverged.

The first consensus strategy was a conservative, lowest-common-ancestor approach. When methods disagreed at fine-grained leaves, votes were propagated to leaf nodes, and the algorithm searched for the lowest ancestor common to all highest-vote leaves. This rule prioritises specificity only when the evidence is concordant across methods; otherwise, it returns a higher-level parent rather than forcing a potentially spurious fine label.

The second strategy was a weighted vote propagation model. We constructed a sparse vote matrix over the hierarchy nodes and propagated votes in two directions. Downstream propagation from each predicted node to its descendants captured support for finer resolutions. Upstream propagation (half the vote at each node) to ancestors attenuated support at each step, encoding the intuition that distant ancestors should not dominate the decision. A down-weighting of its self-vote (90% of the original vote) encouraged the selection of a true common ancestor. The final consensus was the unique node with maximal combined support; in the presence of ties at the maximum, the label was left missing rather than arbitrarily breaking the tie.

To obtain a final ensemble label suitable for downstream analyses, we combined the data-driven consensus from Azimuth, Blueprint, and Monaco with the original CELLxGENE annotation. If the ensemble result and CELLxGENE both indicated non-immune, we retained the original unified label; if the ensemble result indicated non-immune but CELLxGENE indicated an immune type, we assigned “other” to avoid misclassification; otherwise, we kept the ensemble result. If either approach failed to type the cell, we used the other. Finally, for cell types that are only captured in the original annotation, such as NKT cells and mast cells, we used the original regardless of what the ensemble approach determined; however, for NKT cells specifically, we imposed the restriction that the ensemble approach should call these cells as either NK or T cells.

The immune hierarchy used for consensus was read from an immune tree and converted to a directed graph. The immune tree was heuristically adapted and modified from the classical model of the immune cell differentiation^60^. We validated that all harmonised labels were present as nodes, setting invalid entries to missing. Graph distances in the parent and child directions supported the vote propagation and ancestor search procedures. When summarising the depth of annotation, we collapsed labels into fixed hierarchy levels by mapping each node to its ancestor at the requested depth and setting internal nodes that did not match the target depth to missing values to avoid unintended matches. An alternative Human Cell Atlas Ontology (HCAO) immune graph was built semi-programmatically (Supplementary Methods and Supplementary Figure S12) for users seeking finer annotation resolution and more programmatic integration with additional references.

### Pseudobulk

We provide a pseudobulk representation of the cellNexus single-cell resource. Pseudobulk were calculated using the HPCell pipeline (Github: MangiolaLaboratory/HPCell). The process included filtering for high-quality cells (i.e., non-empty droplets, singlets, and alive) and aggregating single-cell gene abundances by sample identifier (sample_id) and ensemble cell typing (cell_type_unified_ensemble), using scater^52^.

### Pseudobulk differential expression analysis of memory B cells

To showcase the effect of improved annotation resolution on integrative analysis, we conducted a pseudobulk differential expression analysis comparing memory B cells in adipose tissue and those in blood. We selected these two tissue groups because there are expectations on what genes may be enriched based on circulation and/or residency. Using the original CELLxGENE annotation, we first selected all B memory cells that came from adipose tissue. Then we selected all blood B memory cells that came from the same dataset as the adipose tissue B memory cells to minimise batch effects. We aggregated the single-cell data of the selected cells as described in the previous section, with the exception that we aggregated by donor instead of sample, and yielded 2 pseudobulk samples for adipose tissue and 12 pseudobulk samples for blood. We filtered the abundant features using the R package tidybulk^53^. We conducted differential expression analysis using the R package variancePartition^61^, with fixed effects of tissue group and sex, and, where applicable, random effects of dataset and donor. We repeated the same analysis using the cellNexus ensemble annotation, yielding 16 adipose tissue samples and 12 blood samples. To assess the effect of annotation resolution improvement, we identified 10 genes that may be enriched in adipose tissue (CXCR4, ITGA4, ITGB1, CD44, BCL2, HLA-DRA, CD69, CD74, NFKBIA, and TNFAIP3) and examined whether they were significantly upregulated in adipose tissue.

### Application programming interface (API)

We developed cellNexus as an R interface that enables reproducible, programmatic access to a harmonised copy of CELLxGENE using a simple query-then-retrieve paradigm. We distribute metadata as Parquet on the ARDC Nectar object store. It is loaded lazily via DuckDB, allowing users to filter with familiar data verbs while queries are pushed down efficiently without materialising large tables. On first use, the backend synchronises any required cloud files to a versioned local cache organised by atlas, aggregation level (single cell and pseudobulk), and assay (counts, counts per million and SCT-normalised^10^), reporting expected download sizes and performing exception-safe, resumable transfers. SCT normalisation was performed (using SCTransform v2^10^) at the sample level for parallelisation, while enforcing a median common scale across the whole resource (scale_factor=2,186). Counts are stored as H5AD and read in HDF5-backed mode to limit memory usage; when multiple files are combined, we reconcile genes via set intersection and bind the matrices while deriving coherent column-level metadata from the filtered table (with safeguards for duplicate identifiers and single-column edge cases). The API returns standard Bioconductor containers (SingleCellExperiment or SummarizedExperiment) to maximise interoperability, with optional conversion to Seurat; higher-level aggregations are supported through principled grouping, including sample-by-cell-type pseudobulk. Beyond the harmonised core, we expose unharmonised, dataset-specific fields as on-demand sidecars and permit integration of user-supplied Parquet or SingleCellExperiment objects through the same validation, caching and directory schema. Throughout, the system emphasises transparency (informative progress messages), safety (input validation, file integrity checks) and portability (cellChat-based, URL-addressable resources), enabling scalable, reproducible single-cell retrieval across large, heterogeneous atlases.

cellNexus provides a simple query-then-retrieve workflow that couples a lazy metadata layer with transparent cloud synchronisation and cache-backed assembly of analysis objects. The entry point, get_metadata(), downloads versioned Parquet metadata from the ARDC Nectar object store to the user cache (get_default_cache_dir()), opens them in DuckDB as lazy dplyr tables with pushdown filtering, and memoises connections by hashing the file set; it also supports mixing cloud_metadata and local_metadata. Filtered rows are then passed to retrieval functions: get_single_cell_experiment() validates required identifiers, synchronises missing H5AD count files into a versioned directory hierarchy by atlas, aggregation, and assay, reads them HDF5-backed, reconciles genes by intersection when needed, and composes a unified SingleCellExperiment with requested assays (counts, counts per million and SCT normalised^10^) and optional features subsetting. Higher-level aggregations follow the same principles: get_pseudobulk() returns a SummarizedExperiment aggregated by sample and harmonised cell type. For interoperability, get_seurat() converts retrieved experiments to Seurat objects without reloading data. Across endpoints, the backend emphasises principled design choices, lazy evaluation for scalability, deterministic cache layout for portability, explicit validation and informative progress/warnings for transparency, and standard Bioconductor/Seurat containers for downstream reproducibility.

### Cell communication analyses

We modelled age-related changes in intercellular communication by analysing ligand-receptor pathway activity across diverse human tissues (CellChat^39^). Communication strength was summarised for each cell-type pair and sample using pseudobulk estimates of log-transformed pathway probabilities, derived from a structured relational database of interaction metrics. Sample-level metadata, including age in days, sex, tissue (grouped into macrocategories; Supplementary Material), donor, disease state (grouped into macrocategories; Supplementary Material), self-reported ethnicity (Figure 4A), and assay type (grouped into macrocategories; Supplementary Material), were harmonised before modelling. We restricted cell-cell pairs to biologically plausible combinations, excluding ambiguous cell labels and retaining immune-immune interactions and immune-non-immune crosstalk in both directions. We performed analyses only for combinations in which at least 100 unique donors supported each (source, target, pathway) combination.

Age was encoded using ordered categorical bins corresponding to Infancy (<1 year old), Childhood (1-10 years old), Adolescence (11-20 years old), Young Adulthood (21-40 years old), Middle Age (41-60 years old), and Senior decades (>60 years old). For each cell-cell-pathway combination, we fit a Bayesian mixed-effect model (Gaussian family) in which log-transformed pathway probabilities were regressed on age bin, sex, and their interaction. The linear model is defined by ∼ age + sex + ethnicity_group + technology_group + disease_group + (1 | dataset) + (age + sex | tissue_group). Using brms^27^ (version 2.23.0) with the Hamiltonian Monte Carlo algorithm, we modelled dispersion as an intercept-only submodel. Priors were weakly informative and adapted to the scale of the interaction weights, with Student-t priors for intercepts and fixed effects, exponential priors for standard deviations, and LKJ priors for correlation matrices.

To test for monotonic shifts in pathway usage with age without assuming linearity, we considered all possible binary partitions of the ordered age bins and compared the mean pathway probability in older versus younger groups. Directional hypotheses were framed with an indifference region (τ = 0.2) on the response scale, evaluating whether the posterior support favoured increased or decreased communication strength beyond this threshold. We calculated posterior probabilities for the positive and negative directions and combined them into a single absolute-direction measure. These tests were carried out both at the population level and with random tissue effects included. All tests are double-sided.

To account for multiple testing across pathways, we applied a Bayesian analogue of the false discovery rate (BFDR), ranking tests by the strength of their posterior support and estimating the expected number of false discoveries among the top-k results. These statistics provided are q-value analogues. To fairly compare pathway shifts across contexts with varying sample sizes, we adjusted pathway counts by regressing their log-counts on sample size and using the residuals as normalised indicators of pathway burden.

Pathway usage profiles were summarised across tissues and cell types using a combination of adjusted counts, pyramidal bar plots, and volcano-style effect-probability displays. For focal pathways, we reconstructed per-sample adjusted values by adding posterior residuals to fixed-effect predictions, with random effects suppressed, and retaining the residuals on the response scale. We then visualised tissue-stratified posterior summaries to illustrate how communication strength is modulated across age bins.

To contextualise macrophage-muscle communication axes, particularly involving the LIF-LIFR pathway, we re-analysed single-cell RNA-seq data from relevant tissues. We filtered cells for quality, normalised them, and embedded them using PCA and UMAP after batch integration. Expression of candidate ligands and receptors was visualised across the cell-state embedding and stratified by coarse age groups to reveal shifts in expression and localisation patterns. Expression maps confirmed coordinated expression of ligand and receptor components in macrophage and muscle-like clusters, respectively.

Visual summaries aggregated pathway-level signals across cell types and functional categories. We grouped ligand-receptor pairs into mechanistic classes (e.g., maintenance, inflammatory, dual-role). We summarised their direction and effect sizes across tissues to characterise the overall ageing trajectory of immune signalling. These summaries were arranged into multi-panel figures that combined signed-count overviews, partner-specific pyramids, pathway-level volcano plots, tissue-specific error bars, and ligand-receptor expression maps overlaid on single-cell embeddings.

The entire analysis was orchestrated using a reproducible workflow in R, encoded as a targets pipeline and executed on a Slurm cluster using an elastic pool of memory-scaled job controllers. Data preparation was optimised using query pushdown in DuckDB, and all intermediate outputs were serialised efficiently to support reproducibility. Key computational steps, including model fitting, hypothesis testing, and plotting, were modularised and versioned, allowing rapid iteration and robust downstream summaries across diverse tissue-cell-pathway contexts.

### Miscellaneous tools

cellNexus and this manuscript was possible thanks to the following software: AnnotationDbi 1.72.0^62^; BiocGenerics data containers ^63^; Azimuth 0.5.0 ^35^; biomaRt 2.66.0^64^; Celldex 1.20.0^38^; CellChat version 2.2.0^39^ tidyverse^49^; DropletUtils 1.30.0 ^65^; edgeR 4.8.0 ^66^; Ensembldb 2.34.0 ^67^; scDblFinder 1.24.0 ^21^; Seurat 5.3.1 ^68^; SingleR 2.12.0 ^38^; Targets 1.11.1^50^; tidybulk 2.1.1^53^; tidyomics 1.6.0^54^; tidyseurat 0.8.7^69^; tidyHeatmap ^70^, igraph^71^.

